# Does temperature change worsen or mitigate the effect of malaria infection on erythrocyte deformability?

**DOI:** 10.1101/136796

**Authors:** A.S. Ademiloye, L.W. Zhang, K.M. Liew

## INTRODUCTION

This paper attempts to investigate whether temperature change worsen or mitigate the effect of *Plasmodium falciparum* (*P. falciparum*) infection and maturation on the deformability of malaria-infected red blood cells (iRBCs) or erythrocytes. *P. falciparum* infection is the most lethal malaria parasitic infection affecting humans [1]. Upon infection, the merozoites enter the bloodstream and develop through the ring (*Pf*-rRBC), trophozoite (*Pf*-tRBC) and schizont (*Pf*-tRBC) stages, which is usually accompanied by significant changes in body temperature. Hence, this study aims to draw a connection between temperature change and erythrocyte deformability upon malaria infection.

## METHODS

We employed a newly proposed 3D multiscale semi-analytical framework [2] to predict the elastic properties of the iRBC membrane. Summarily, the atomistic scale strain energy density function (*W*) of the iRBC membrane is obtained from the summation of membrane’s in-plane energy and bending energy that are computed using the cytoskeletal microstructure parameters such as spectrin link length (*L*), persistence length (*p*) and maximum contour length (*L*_max_). Assume that a selected membrane representative cell is subjected to an area-preserving shear deformation; the membrane elastic properties can be obtained by minimizing the *W* of the representative cell using Newton method with respect to four geometric parameters defined in a 3D space as detailed in Refs. [2–5].

## RESULTS AND DISCUSSION

Using the set of microstructure parameters in Table 1, the Young’s modulus (*E*) and shear modulus (*µ*) of the iRBC membrane are computed at various temperature values.

**Table 1.**
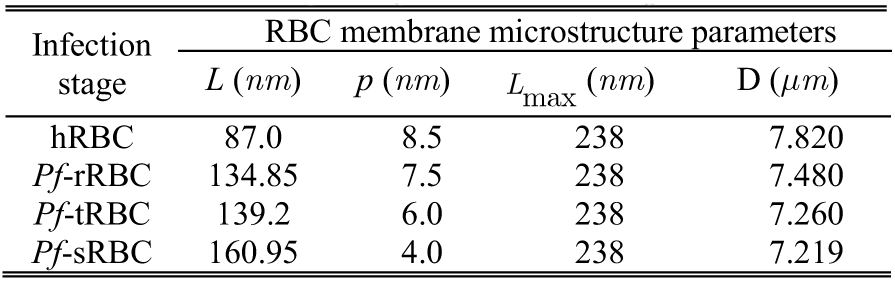
Values of erythrocyte microstructure parameters.

The results obtained from this study reveal that a multifold increase in elastic properties occurs as infection progresses. In addition, membrane stiffness increases as temperature rises (see Table 2). It is concluded that the effect of malaria infection worsen with increase in temperature, resulting to significant reduction in erythrocyte deformability.

**Table 2.**
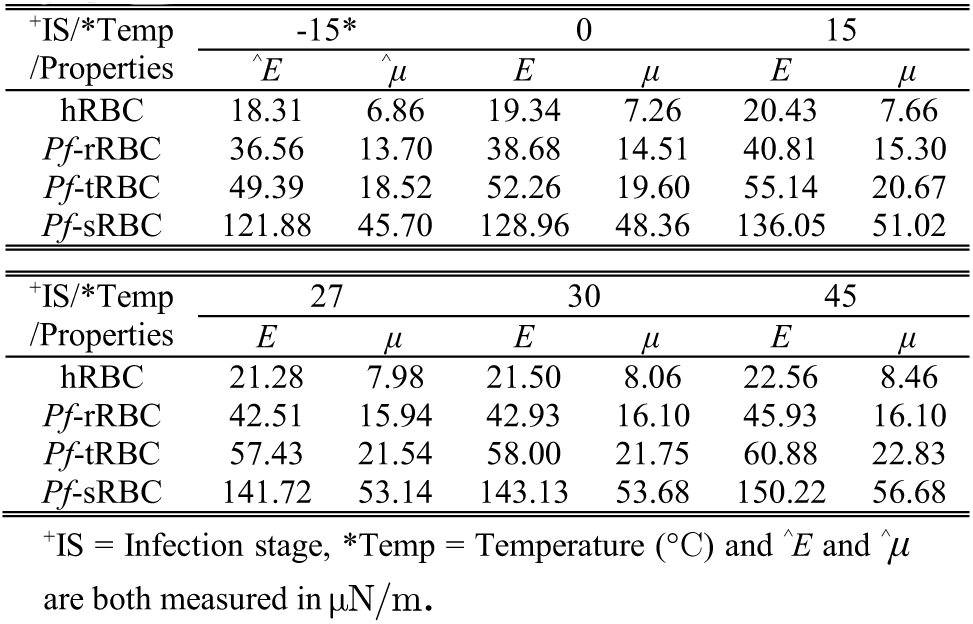
Effect of temperature on iRBC membrane elastic properties.

| +IS/*Temp /Properties | -15* |  | 0 |  | 15 |  |
| --- | --- | --- | --- | --- | --- | --- |
| | $\hat{E}$ | $\hat{\mu}$ | $E$ | $\mu$ | $E$ | $\mu$ |
| hRBC | 18.31 | 6.86 | 19.34 | 7.26 | 20.43 | 7.66 |
| *Pf*-rRBC | 36.56 | 13.70 | 38.68 | 14.51 | 40.81 | 15.30 |
| *Pf*-tRBC | 49.39 | 18.52 | 52.26 | 19.60 | 55.14 | 20.67 |
| *Pf*-sRBC | 121.88 | 45.70 | 128.96 | 48.36 | 136.05 | 51.02 |

| +IS/*Temp /Properties | 27 |  | 30 |  | 45 |  |
| --- | --- | --- | --- | --- | --- | --- |
| | $E$ | $\mu$ | $E$ | $\mu$ | $E$ | $\mu$ |
| hRBC | 21.28 | 7.98 | 21.50 | 8.06 | 22.56 | 8.46 |
| *Pf*-rRBC | 42.51 | 15.94 | 42.93 | 16.10 | 45.93 | 16.10 |
| *Pf*-tRBC | 57.43 | 21.54 | 58.00 | 21.75 | 60.88 | 22.83 |
| *Pf*-sRBC | 141.72 | 53.14 | 143.13 | 53.68 | 150.22 | 56.68 |

+IS = Infection stage, \*Temp = Temperature ( $^{\circ}$ C) and $\hat{E}$ and $\hat{\mu}$ are both measured in $\mu$ N/m.

## ACKNOWLEDGEMENT

The study was financially supported by the RGC, HKSAR (Project No. 9042047, CityU 11208914) and NSFC, China (Grant No. 11402142 and Grant No. 51378448).

